# Initiation phase cellular reprogramming ameliorates DNA damage in the ERCC1 mouse model of premature aging

**DOI:** 10.1101/2023.05.12.540500

**Authors:** Patrick Treat Paine, Cheyenne Rechsteiner, Francesco Morandini, Gabriela Desdin Mico, Calida Mrabti, Alberto Parras, Amin Haghani, Robert Brooke, Steve Horvath, Andrei Seluanov, Vera Gorbunova, Alejandro Ocampo

## Abstract

Unlike aged somatic cells, which exhibit a decline in molecular fidelity and eventually reach a state of replicative senescence, pluripotent stem cells can indefinitely replenish themselves while retaining full homeostatic capacity. The conferment of beneficial-pluripotency related traits via in vivo partial cellular reprogramming (IVPR) significantly extends lifespan and restores aging phenotypes in mouse models. Although the phases of cellular reprogramming are well characterized, details of the rejuvenation processes are poorly defined. To understand whether epigenetic reprogramming can ameliorate DNA damage, we created reprogrammable accelerated aging mouse model with an ERCC1 mutation. Importantly, using enhanced partial reprogramming by combining small molecules with the Yamanaka factors, we observed potent reversion of DNA damage, significant upregulation of multiple DNA damage repair processes, and restoration of the epigenetic clock. In addition, we present evidence that pharmacological inhibition of ALK5 and ALK2 receptors in TGFb pathway is able to phenocopy some benefits including epigenetic clock restoration suggesting a role in the mechanism of rejuvenation by partial reprogramming.

## Introduction

The phenomenon of aging is directly linked to a decline in cellular repair functions with an associated increase in aging phenotypes including genomic instability^1–3^. DNA damage events due to radiation, reactive oxygen species, chemicals, or replication errors can overwhelm DNA repair functions and are proposed as a causative factor for epigenetic dysregulation and a key contributor to age associated pathologies^4–8^. In this line, Ercc1^Δ/−^ progeroid mice, harboring a single truncated Ercc1 allele required for nucleotide excision repair (NER), interstrand crosslink repair (ICL), and homologous repair (HR) display increased DNA damage and a broad spectrum of aging phenotypes including senescence, neurodegeneration, multi-morbidity, and a shortened lifespan^9–13^. Interestingly, a recent comparative lifespan analysis of 18 wildtype rodent species indicates DNA double-strand break repair was more efficient in long-lived species^14^. For these reasons, DNA repair mutant Ercc1^Δ/−^ mice are an ideal alternative for investigating aging interventions and DNA repair-related mechanisms of rejuvenation in particular^9, 15, 16^.

Cellular reprogramming can be defined as the conversion of a somatic cell to pluripotency and can be induced via the forced expression of four defined transcription factors; Oct4, Sox2, Klf4, and c-Myc (OSKM)^17^. The resulting induced pluripotent stem cells (iPSCs) exhibit a dedifferentiated cell identity similar to embryonic stem cells (ESCs) along with a restoration of aged phenotypes^18, 19^. Recently, in vivo partial reprogramming (IVPR), following short-term cyclic expression of OSKM, has emerged as a novel therapeutic strategy for the treatment of age-related diseases (ARDs)^20–22^. Specifically, partial cellular reprogramming has produced improvements to lifespan, hallmarks of aging, the epigenetic clock, and tissue regeneration in progeroid and wildtype mouse models in vivo^6, 23–34^. Multiple in vitro time course studies have demonstrated reprogramming initially proceeds in a multifactorial manner involving extensive epigenetic remodeling, cell cycle induction with a shorted G1 phase, mesenchymal to epithelial transition due to TGFb inhibition, and BMP induction, while undergoing a major reset to the proteome and transcriptome^35–45^. Importantly, in what manner and to what extent these early alterations to molecular and cellular processes via IVPR counter and reverse aging drivers is currently unknown.

To address this limitation and gain a deeper understanding of the mechanisms involved in age amelioration by OSKM induction, we have chosen to investigate the effects of partial reprogramming in the Ercc1^Δ/−^ DNA damage model of accelerated aging. Using an in vitro time course capable of dissecting time dependent processes, we characterized the early events of reprogramming in a novel reprogrammable Ercc1^Δ/−^ accelerated aging mouse model. Most importantly, we observed a significant reduction in DNA damage beginning at 2 days of reprogramming demonstrating reversal of a key hallmark of aging. At the same time RNA seq analysis shows a significant upregulation of nearly every major DNA repair pathway while DNA methylation clock analysis shows a reversal of epigenetic age. Interestingly, improvements were more robust in the Ercc1 DNA damage model versus the wildtype cells, as would be expected from an improvement to homeostatic capacity. Lastly, small molecule inhibition of the TGFb pathway was sufficient to phenocopy some rejuvenating aspects of reprogramming including a decrease in nuclear size, decreased yH2AX, and restoration of the epigenetic clock.

## Results

### Induced reprogramming in a mouse model of accelerated aging decreases DNA damage

OSKM-mediated partial cellular reprogramming has been shown to mitigate genomic instability and epigenetic alterations in wildtype and Lamin A mutant mouse fibroblasts as well as in vitro aged nucleus pulposus cells^23, 31^. On other hand, it is currently unknown whether this restoration would occur in an Ercc1 DNA damage model of accelerated aging with defects in DNA repair. To address this question, we developed a novel doxycycline-inducible mouse model of reprogrammable aging (Ercc1^Δ/−^ 4Fj^+/−^ rtTA^+/−^) containing a single truncated Ercc1 allele, Col1a1-tetO-OKSM polycistronic transgene, and the ROSA26-M2-rtTA allele (Fig. 1a).

**Figure 1:**
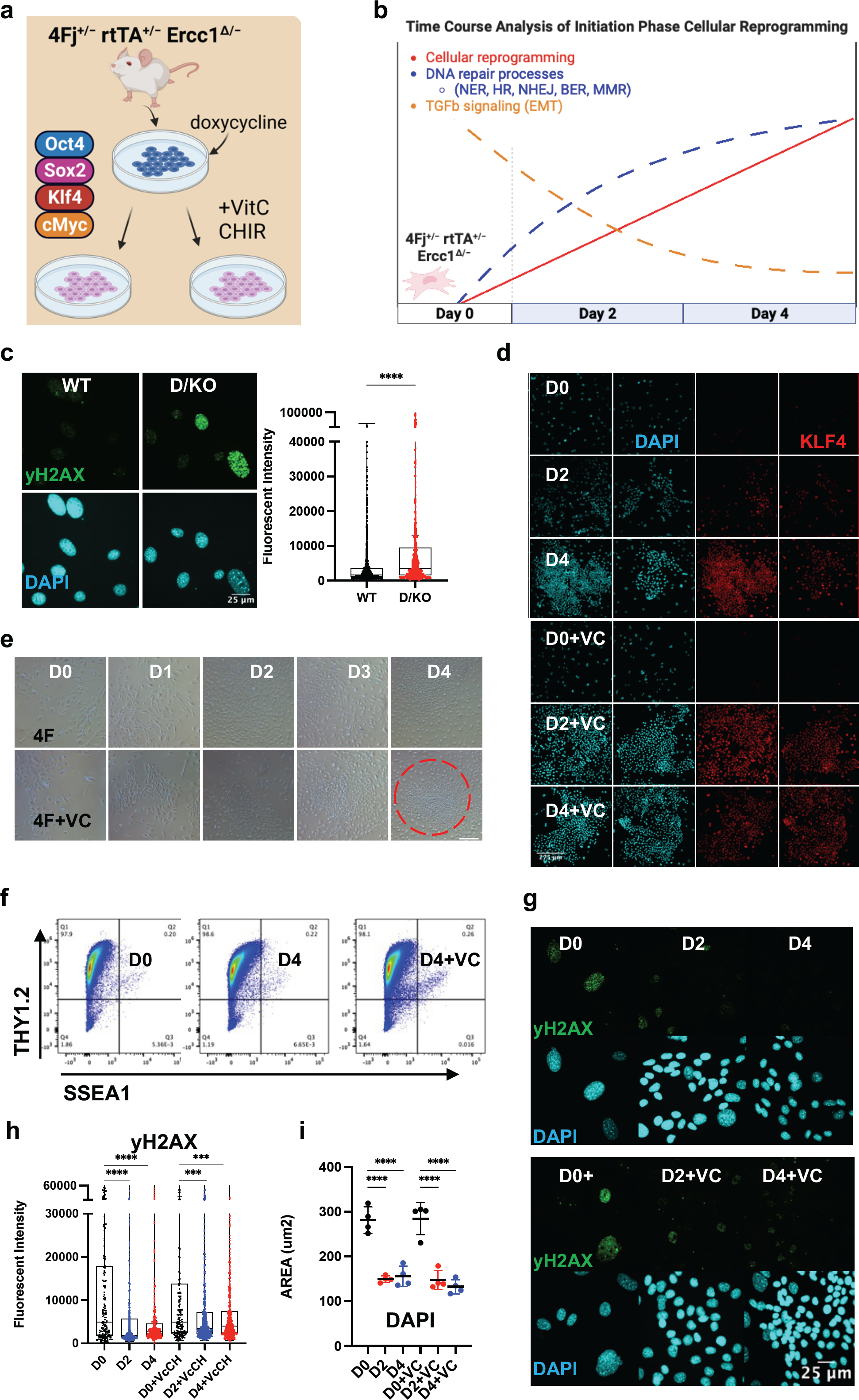
Initiation phase reprogramming in 4Fj^+/−^ rtTA^+/−^ Ercc1^Δ/−^ accelerated aging model promotes DNA damage repair. **(a)** Illustration of doxycycline inducible cellular reprogramming Ercc1 mutant fibroblasts (Fb) alone or enhanced with small molecules. **(b)** Schematic of initiation phase timecourse analysis setup with observed changes to DNA repair and TGFb signaling. **(c)** Representative Immunofluorescence (IF) and quantification of yH2AX in fibroblasts from wildtype (WT) and Ercc1^Δ/−^ (D/KO) mice, imaged with Nikon laser confocal spinning disc, according to Mann-Whitney test, **** p<0.0001. **(d)** IF of KLF4 and DAPI in 4Fj^+/−^ rtTA^+/−^ Ercc1^Δ/−^ Fbs after 2 and 4 days doxycycline induction with or without VC at 100x **(e)** Time course analysis of 4F and 4F.D/KO Fb during 4 days of dox induction +/-VC, brightfield (BF) images at 4x. **(f)** FACS analysis of reprogramming FBs at day 0 and day 4, stained with Thy1.2 and SSEA1. **(g)** IF of yH2AX and DAPI during time course analysis of 4F.D/KO Fb +/-VC. **(h)** IF quantification of yH2AX levels during timecourse shows decrease levels at day 2 and day 4 according to one way ANOVA, *** p<0.001. **(i)** Quantification of DAPI area during timecourse shows decrease in size at day 2 and day 4 based on mean values of 4 experiments according to one way ANOVA, **** p<0.0001.

To first verify the presence of DNA damage in our reprogrammable progeria model, adult tail tip fibroblasts (FBs) were isolated from the 4Fj Ercc1^Δ/−^ (D/KO) and 4Fj Ercc1^+/+^ (WT) mice, stained for the DNA damage marker γH2AX, and imaged with confocal laser microscopy. As expected, a significant increase in yH2AX fluorescence was observed in the D/KO fibroblasts compared to the WT (Fig. 1c, s1a). In addition, this accelerated aging cell model also displayed a significant increase in nuclear area in these samples, based on nuclear staining with DAPI (Fig. S1b). This experiment confirmed the presence of a DNA repair defect in our novel OSKM inducible mouse model of aging along with an enlarged nucleus compared to WT (Fig. S1a,b).

In vitro cellular reprogramming time course experiments have previously been used to identify three phases of reprogramming including initiation, maturation, and stabilization^46, 47^. Initiation phase proceeds via alterations to several key biological processes including proliferation, chromatin modification, DNA damage repair, mesenchymal to epithelial transition (MET), and RNA processing^33, 41–45, 48–50^. To gain insight into partial reprogramming processes with the capacity to restore homeostatic function in a DNA damage model, a time course analysis of the initiation phase of reprogramming was performed (Fig. 1b). At the same time, the induction of pluripotency in somatic cells following OSKM expression is a stochastic, asynchronous, and inefficient process often taking weeks with a low percentage of iPSC colonies^51, 52^. To facilitate an accelerated reprogramming process and enhance the ability to delineate reprogramming-induced aging phenotypes in bulk cultures a previously identified small molecule combination, Vitamin C (V) and CHIR-99021 (C), was selected^53–55^. Alone, Vitamin C is able lower the epigenetic barrier of pluripotent gene expression due to its function as a histone demethylase and Tet enzyme cofactor while CHIR activates the Wnt pathway and promotes glycolysis via GSK3-beta inhibition^56–58^.

To confirm the induction of cellular reprogramming following expression of OSKM in our reprogrammable DNA damage model, doxycycline (2ug/mL) was added to 4F Ercc1 cells in culture alone or combined with VC and imaged with brightfield microscopy at 4x during days 0 – 4 (Fig. 1a,b). A clear shift in cell morphology with a cobblestone appearance associated with MET is present after 2 days, more so after 4 days, and most strongly in the enhanced D4+VC group (Fig. 1e). Expression of the Klf4 and Sox2 transcription factors were observed only after doxycycline induction and most strongly at d4+VC but never day 0 based on immunofluorescence (IF) laser confocal imaging (Fig. 1d, S1c). These cells retained the fibroblast identity marker Thy1.2 at D4 with or without VC, as shown by flow cytometry, indicating this mouse model and experimental setup successfully represent initiation phase partial reprogramming without dedifferentiation or entrance into a pluripotent state (Fig. 1g, s1d).

In regards to the effects of cellular reprogramming on yH2AX in this DNA damage model, we first used flow cytometry. Specifically, two days of doxycycline induced reprogramming in the D/KO was sufficient to decrease median yH2AX fluorescent intensity per cell by 38% (Fig. S1e). Next, using IF and confocal imaging, we observed a significant decrease in mean yH2AX level per cell after 2 and 4 days of reprogramming (Fig. 1g, h, S1j). Similarly, enhanced reprogramming with VC also restored this DNA damage marker significantly after 2 and 4 days of OSKM induction (Fig. 1g, h, S1j). These D/KO cells also displayed a significant reduction in nuclear size during reprogramming after induction with doxycycline for 2 or 4 days (Fig. 1i). When this experiment was repeated on OSKM inducible WT cells without the DNA repair defect, it also showed a significant decrease in yH2AX signal and decrease in nuclear area (Fig. S1f). Thus, initiation phase cellular reprogramming is sufficient to ameliorate a key driver of aging even in mutant cells defective for DNA repair responsible for NER, ICL, and HR.

### The DNA methylation clock is restored in Ercc1^Δ/−^ following short term reprogramming

Recent evidence from our lab indicates that D/KO FBs display accelerated aging based on the DNA methylation clock and are therefore a good *in vitro* model to investigate aging mechanisms and interventions^59^. Furthermore, cellular reprogramming has been observed to reverse epigenetic age in mouse and human cell types but the effects on an accelerated aging model with a DNA repair defect are unknown^60, 61^. In order to investigate the effect of reprogramming on the DNA methylation clock, we applied the DNA Methyl Age Skin Final clock to all samples and timepoints. Strikingly, enhanced reprogramming with VC for 4 days in the D/KO produced a significant restoration to the DNA methylation clock with the top responder showing a 54% decrease in epigenetic age (Fig. 2a). At the same time, there was a trend towards epigenetic clock restoration after 2 and 4 days of reprogramming although not significant, perhaps due to less efficient reprogramming (Fig. 2a). Significant restoration to the DNAm clock only occurred at day 2 in WT FBs (Fig. s2a). In addition, variability was greater in the WT samples perhaps due to the stochastic nature of reprogramming and the early time point chosen for analysis, or the lack of accelerate aging phenotype (Fig. s2a). The robust reversal of the DNAm clock in the D/KO FBs indicates enhanced reprogramming is capable of cellular rejuvenation in an aging model with defective DNA repair.

**Figure 2:**
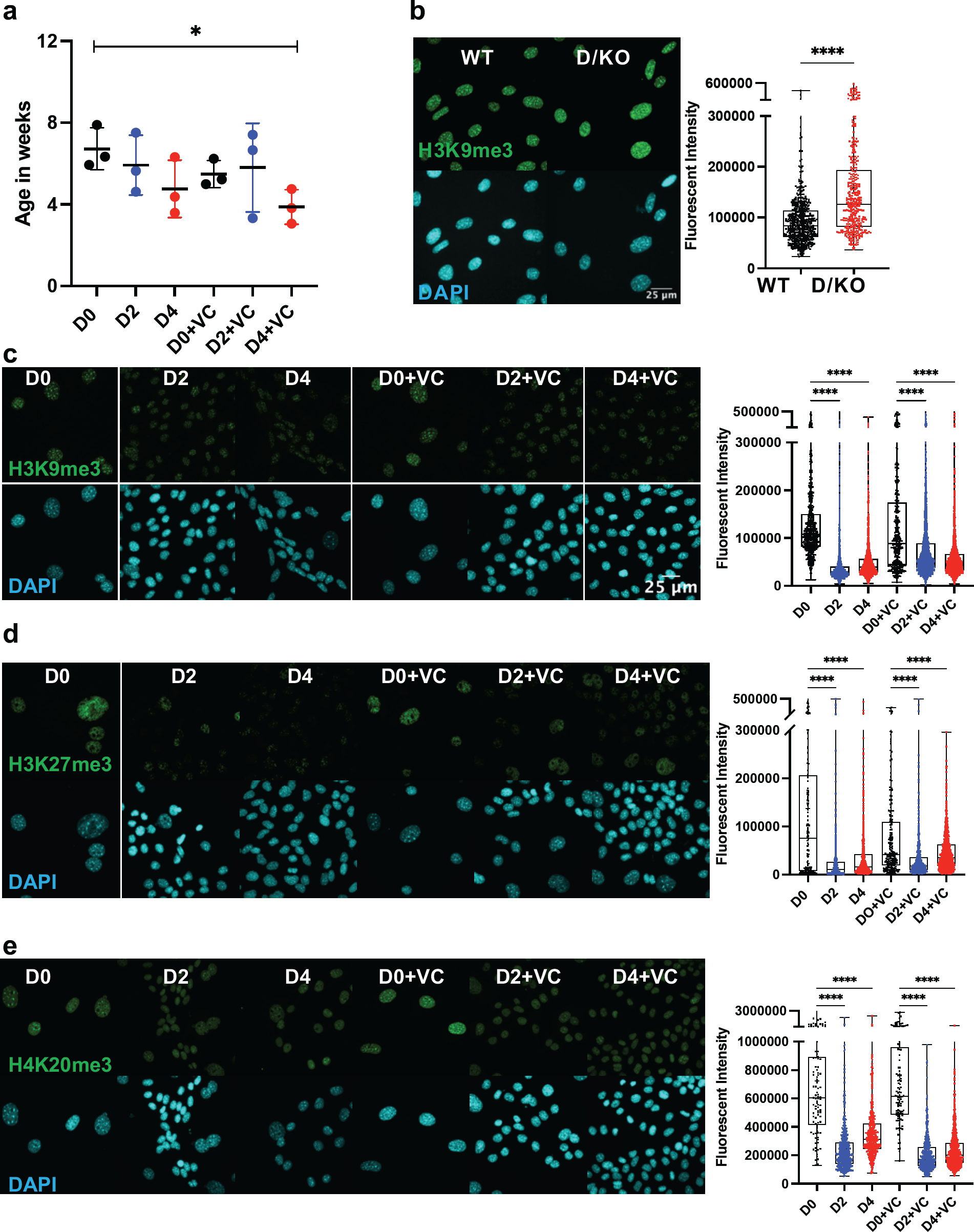
DNA methylation clock is restored in Ercc1^Δ/−^ following short term reprogramming. **(a)** DNA methylation clock (DNA Methylation Age Skin Final clock) analysis of cellular Ercc1 mutant Fb during time course, n = 3, * p<0.05 according to unpaired t-test. **(b)** IF and quantification of H3K9me3 in fibroblasts from WT and D/KO mice, imaged with Nikon laser confocal spinning disc, according to Mann-Whitney test, **** p<0.0001. **(c)** IF and quantification of H3K9me3 and DAPI during time course in 4F.D/KO Fbs after 2 and 4 days doxycycline induction with or without VC at 100x according to Kruskal-Wallis test, * p<0.0001 **(d)** IF and quantification of H3K27me3 and DAPI during time course in 4F.D/KO Fbs after 2 and 4 days doxycycline induction with or without VC at 100x according to Kruskal-Wallis test, * p<0.0001 **(e)** IF and quantification of H4K20me3 and DAPI during time course in 4F.D/KO Fbs after 2 and 4 days doxycycline induction with or without VC at 100x according to Kruskal-Wallis test, * p<0.0001.

Next, as chromatin remodeling and reversal of the DNA methylation clock are observed to coincide during cellular reprogramming, we sought to understand the epigenetic alterations that occur following reprogramming in this Ercc1 model^48, 61^. First, a significant increase in H3K9me3 was observed in the D/KO fibroblasts compared to WT controls based on IF confocal imaging at 100x (Fig. 2b, s2b). A similar increase was observed in the heterochromatin mark H4K20me3 while H3K27me3 remained unchanged under the same conditions (Fig. s2d, e). These results were then confirmed with western capillary analysis (Fig s2c). We hypothesize that this *in vitro* phenotype of chromatin compaction is an adaptation in these cells to protect against chronic elevated DNA damage levels^62, 63^. Interestingly, when reprogramming or enhanced reprogramming was induced for 2 or 4 days, heterochromatin was significantly decreased in this DNA damage model shifting it towards wildtype levels (Fig. 2c, d, e, s2g, h, I, and j). This restoration of heterochromatin levels in D/KO FBs coincides with improvement to reversion of the DNAm clock, in line with previous reports suggesting a mechanistic relationship^6, 23, 32^.

### Homeostatic capacity is significantly upregulated during the initiation phase of cellular reprogramming

To better understand the effects of reprogramming in this DNA damage model, paired-end bulk RNA sequencing and analysis was performed for each treatment and time point. Principal component analysis (PCA) based on relative gene expression values confirmed groups clustered based on the treatment, timepoint, and cell type while displaying a reprogramming trajectory from day 0 to day 4 (Fig. 3a). Interestingly, the enhanced reprogramming at day 2 plus VC was equal to 4 days of reprogramming alone based on overlapping PCA scores (Fig. 3a). At the same time, D/KO cells displayed a larger shift in PC2 score than WT following reprogramming (Fig. 3a). Venn diagrams and associated gene expression heat maps were created based on the normalized transcriptomes from each group and demonstrate a shared gene profile during reprogramming albeit with some unique differences (Fig. 3b, 3c). In particular, over 1400 genes were uniquely upregulated and 992 downregulated in the D/KO Fbs during initiation phase with enhanced reprogramming compared to reprogrammed WT cells (Fig. 3c). The most robust changes were observed following enhanced reprogramming to the D/KO group, indicating a profound and rapid reset to the transcriptome in this DNA damage model (Fig. 3b, 3c). GO term analysis demonstrated a significant upregulation of DNA damage repair pathways in the D/KO enhanced reprogramming group including DNA repair, homologous recombination (HR), non-homologous end joining (NHEJ), base excision repair (BER), mismatch repair (MMR), nucleotide excision repair (NER), and alternative end joining (AltEJ), while interstrand crosslink repair (ICR) showed a trend towards upregulation. (Fig. 3d, S3a). At the same time, reprogramming in WT cells did not produce such broad and robust effects on DNA repair processes (Fig. 3d, S3a). Other notable processes increased with enhanced reprogramming include chromatin organization pathways in D/KO cells (Fig. 3d, S3a). Importantly, there was also a significant downregulation in TGFb receptor signaling and TGFb regulation pathways as well as a decrease in EMT pathways (Fig. 3d, S3a). Gene set enrichment analysis (GSEA) showed that intermediate filament cytoskeleton organization and processes, keratinocyte differentiation and epidermal differentiation were significantly upregulated at all timepoints and models (Fig. S3b). At the same time, Reactome analysis showed that Keratinization and Formation of the Cornified Envelope were also universally upregulated (Fig. S3c). Together, this transcriptomic data provides a mechanistic basis that supports the observed changes to morphological, epigenetic, and DNA damage phenotypes that occur following short-term reprogramming in our DNA damage model. Interestingly, the robust upregulation of major DNA repair processes in the Ercc1 model vs WT suggest a homeostatic process capable of responding to intrinsic molecular and age-related defects that characterized Ercc1 cells.

**Figure 3:**
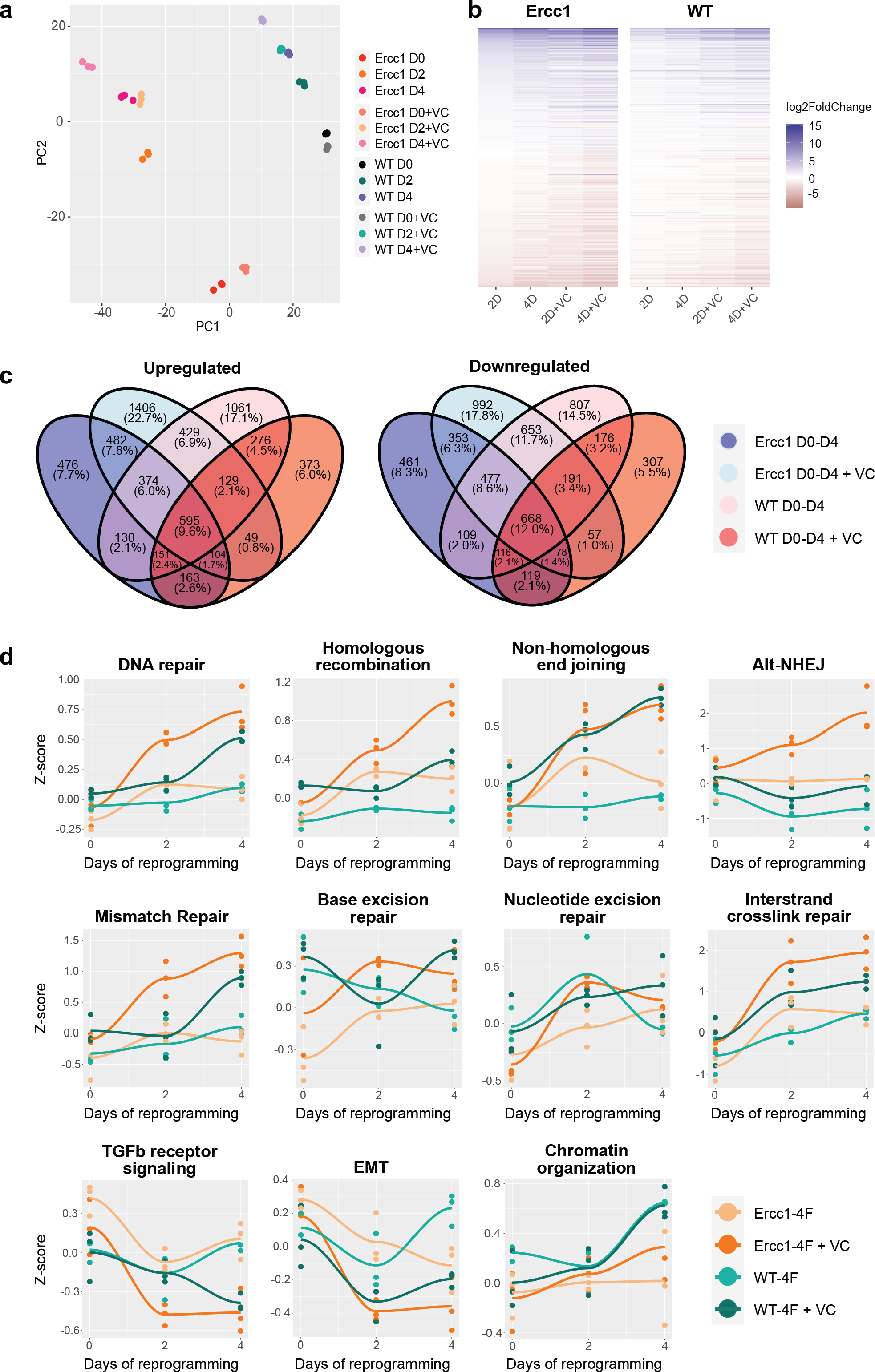
Initation phase reprogramming drives transcriptomic reset and upregulates DNA repair and chromatin organization in Ercc1^Δ/−^. **(a)** Principal component analysis (PCA) of the Ercc1 and WT reprogramming at days 0, 2, and 4 with and without VC enhancement. Principal components 1 and 2 are shown. **(b)** Heat map comparing Ercc1 and WT reprogramming at all timepoints, models, and treatements. Genes with a p-value lower than 1e-18 in at least one condition are shown. **(c)** Venn Diagram showing the overlap of significant DEGs (adjusted p-value < 0.05) in Ercc1 and WT during IP reprogramming with and without VC enhancement. Genes were evaluated using a continuous model over 4 days of reprogramming. **(d)** Reprogramming trajectory visualizations based on median Z-scores of genes from significantly up or down regulated GO pathway including DNA repair, TGFb signaling, EMT, and chromatin organization (n=3).

### TGFb inhibition alone improves DNA damage phenotypes and rejuvenates the DNA methylation clock

The RNA seq analysis showing downregulation of TGFb signaling and associated epithelial to mesenschymal transition pathways is a well-documented early event of cellular reprogramming following OSKM induction^35, 40, 41, 64^. In contrast, upregulation of TGFb signaling is a driver of aging phenotypes including cell degeneration, fibrosis, ROS, inflammation, DNA damage, senescence, and stem cell aging^65–68^. Recently our lab observed that inhibition of the TGFb pathway is able to extend lifespan in C. elegans, supporting its role in aging and longevity^69^. Based on these observations, we asked whether TGFb inhibition alone could impact DNA damage and the DNA methylation clock in our Ercc1 accelerated aging model in a manner similar to reprogramming (Fig. 3a). In this line, the TGFb superfamily consists of over 30 subtypes but for this study we focused on canonical TGFb signaling and the bone morphogenic protein pathway (BMP) as both have been identified to be important for early stage reprogramming ^41, 53, 70^. Specifically, 9 inhibitors that target the TGFb ALK5 receptor and 6 inhibitors that target the BMP ALK2 receptors were screened for 3 days on D/KO fibroblasts. Following a cell viability study to confirm a safe dosage range, a high or low dose of each inhibitor was added to the Ercc1 fibroblasts for 3 days and analyzed for changes to yH2AX using IF and confocal microscopy (Fig. S4a). Surprisingly, all of the ALK5 inhibitors successfully decreased yH2AX levels based on IF, although in some cases it was dose-dependent (Fig. 4b). Similarly, all but one of the ALK2 inhibitors also decreased DNA damage in Ercc1 fibroblasts, although not as effectively as the ALK5 inhibitors, again in a dose dependent manner (Fig. 4c). Subsequently, we repeated these experiments with only the top performers including ALK5 inhibitors Repsox and A83-01, ALK2 inhibitor DMH-1, and dual ALK5 and ALK2 inhibitor Vactosertib. Once again, each of the inhibitors decreased the yH2AX signal of Ercc1 fibroblasts below control levels although the ALK2 inhibitor, DMH-1, was less effective than the three ALK5 inhibitors (Fig. s4b). Interestingly, when nuclear area was calculated based on DAPI staining, DMH-1 was also the only inhibitor not to show a trend towards decreased nuclear size potentially implying an association between nuclear size changes and improvements in DNA repair (Fig. s4). In contrast, other small molecules previously shown to induce pluripotency when applied as a cocktail were unable to decrease yH2AX levels in Ercc1 fibroblasts when applied individually including valproic acid (VPA), CHIR, tranylcypromine (TCP), Forskolin, DZNep, or TTNPB (Fig. s4c) ^71, 72^. When the ALK5 inhibitor, Repsox, was combined with 4 days of reprogramming, no added benefit to yH2AX levels was observed (Fig. S4d). Finally, significant rejuvenation of the DNA methylation clock was observed with 3 of the 4 inhibitors, including Repsox, A83-01, and DMH-1 (Fig. 4d). This data demonstrates that ALK5 and ALK2 inhibition is able reduce DNA damage and restore the DNA methylation clock in this Ercc1 accelerated aging model.

**Figure 4:**
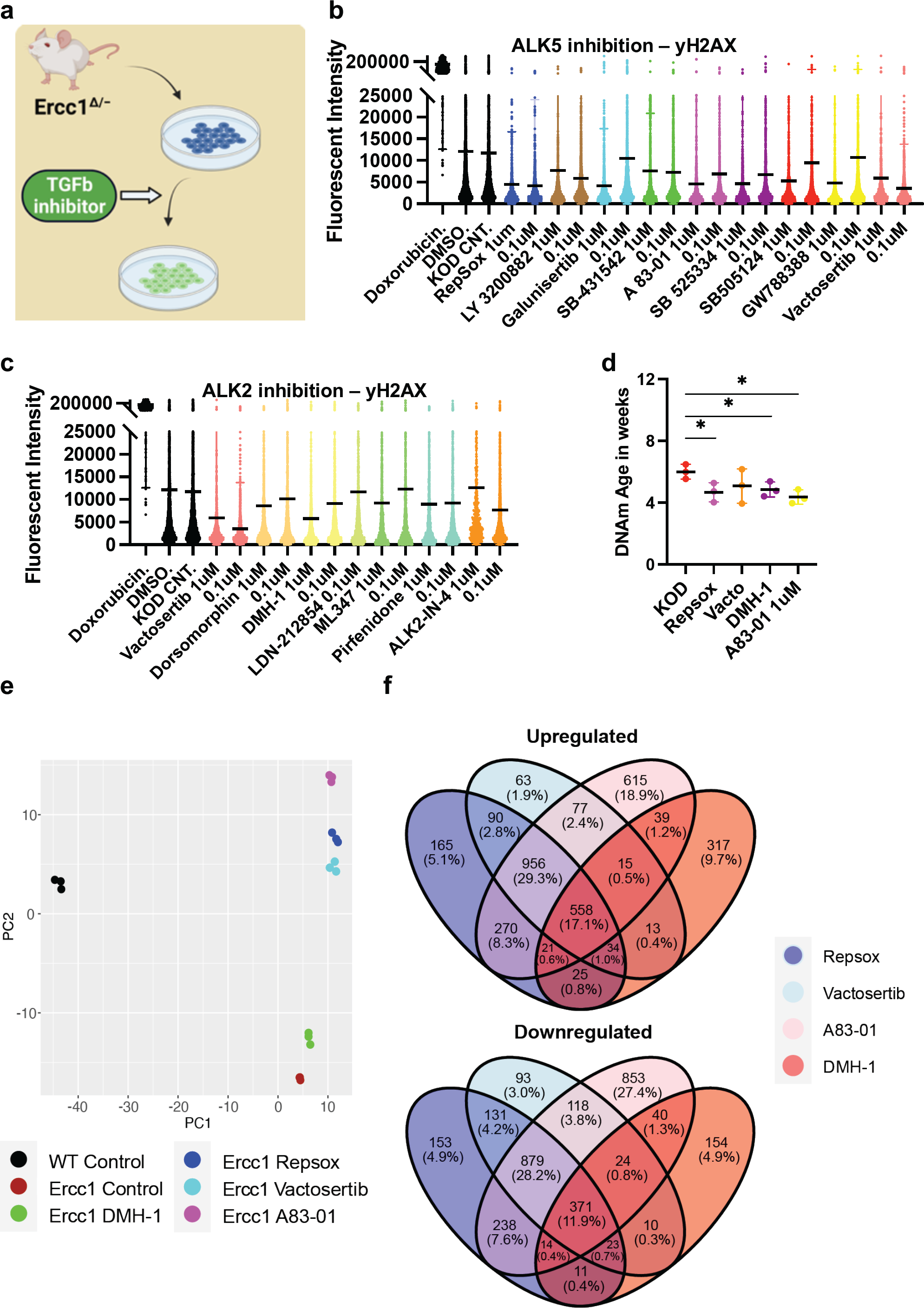
Inhibition of ALK5 or ALK2 receptors improves DNA damage phenotype and resets the DNA methylation clock and transcriptome in Ercc1^Δ/−^. **(a)** Illustration of TGFb inhibitor screen on D/KO Fbs. **(b)** IF quantification of yH2AX mean levels in D/KO Fbs following treatment with ALK5 inhibitors, plated in 96 wells in triplicate and imaged with laser confocal at 40x. **(c)** IF quantification of yH2AX mean levels in D/KO Fbs following treatment with ALK 2 inhibitors, plated in 96 wells in triplicate. **(d)** DNA methylation clock (DNA Methylation Age Skin Final clock) analysis, comparing Repsox (0.1 um), Vactosertib (0.1 um), DMH-1 (1.0 um), and A83-01 (1.0 um) on D/KO Fbs, n = 3, * p<0.05 according to unpaired t-test. **(e)** Principal component analysis (PCA) of Ercc1 cells treated with TGFb inhibitors. Principal components 1 and 2 are shown. **(f)** Venn Diagram showing the overlap of significant genes (adjusted p-value < 0.05) for the TGFb inhibitors.

RNA-seq analysis was next performed in Ercc1 FBs following treatment with these four TGFb inhibitors to better understand the mechanistic basis for improved DNA damage and a rejuvenated DNA methylation clock with ALK2 or ALK5 inhibition. Notably, PCA scores show the three ALK5 inhibitors cluster tightly together compared to the ALK2 inhibitor (Fig. 4e). Interestingly, all 4 inhibitors shifted the D/KO transcriptome along the PC2 axis in the direction of WT cells with the greatest change observed following treatment with A83-01 (Fig. 4e). At the same time, Venn diagrams show 558 genes were upregulated and 371 downregulated by all 4 inhibitors indicating similarity in gene expression profile regardless of target receptor (Fig. 4f). Next, gene set enrichment analysis (GSEA) shows significant decreases to spindle checkpoint signaling with Repsox, Vactosertib, and A83-01 but not with DMH-1 (Fig. S4e). DMH-1 had a unique effect on upregulation of response to virus, response to interferon-gamma, and response to interferon-beta among others based on the GSEA (Fig. S4e). Repsox uniquely induced upregulation to developmental-related gene sets including embryonic hindlimb morphogenesis, hindlimb morphogenesis, and midbrain development (Fig. S4e). Reactome analysis showed that DMH-1 also uniquely increased interferon signalling (Fig. S4e). Interestingly, GO term analysis demonstrated that all four inhibitors significantly downregulated several biological processes including mitotic cell cycle and canonical glycolysis (Fig. S5a). In line with previous publications showing TGFb inhibition disrupts DNA repair processes in cancer cells and HSCs, these four inhibitors also disrupted several DNA repair proceses in our Ercc1 FBs (Fig. S5a)^65, 73–75^. Interesting exceptions include a significant improvement to NER by DMH-1 and a trend towards upregulated ICL repair amongst all (Fig. S5a) Together this data shows ALK5 and ALK2 inhibitors are capable of decreasing the DNA damage marker yH2AX and restoring the DNA methylation clock in Ercc1 fibroblasts while sharing some changes to the transcriptomic profiles.

### TGFb inhibition phenocopies aspects of initiations phase cellular reprogramming

To investigate and compare shared rejuvenation processes between cellular reprogramming and TGFb inhibition, we next evaluated significant transcriptomic changes among the different treatments. Notably, a heatmap of significantly up or down regulated genes among the different treatments indicate a similar profile (Fig. 5c). Specifically, 684 shared DEGs were upregulated and 627 downregulated following treatment with the ALK5 inhibitors Repsox, A83-01, Vactosertib, and reprogramming at day 4 (Fig. 5a). Notably, gene expression changes induced by the 4 inhibitors shared a positive correlation with gene expression changes induced by reprogramming (Fig. 5b). The strongest correlation was observed between the three ALK5 inhibitors and D4 reprogramming (Fig. 5b, S5b). Interestingly, A83-01 was the closest in transcriptomic profile to day 4 reprogramming while Repsox was the closest to D4 enhanced reprogramming (Fig. 5b, S5b). The top GO terms that were significantly downregulated with either reprogramming or the TGFb inhibitors include external encapsulating structure organization, extracellular matrix organization, extracellular structure organization, collagen fibril organization, skeletal system development and protein hydroxylation (Fig. 5d). The most significantly upregulated GO terms shared by both D4 reprogramming and TGFb inhibition include steroid biosynthetic process, regulation. of steroid biosynthetic process, alcohol biosynthetic process, organic hydroxyl compound metabolic process, epidermal cell differentiation, and alcohol metabolic process (Fig. 5d). Reactome analysis of shared processes between reprogramming and Repsox, A83-01, and Vactosertib showed upregulation of complement cascade and downregulation of collagen formation and ECM organization (Fig. 5e).

**Figure 5:**
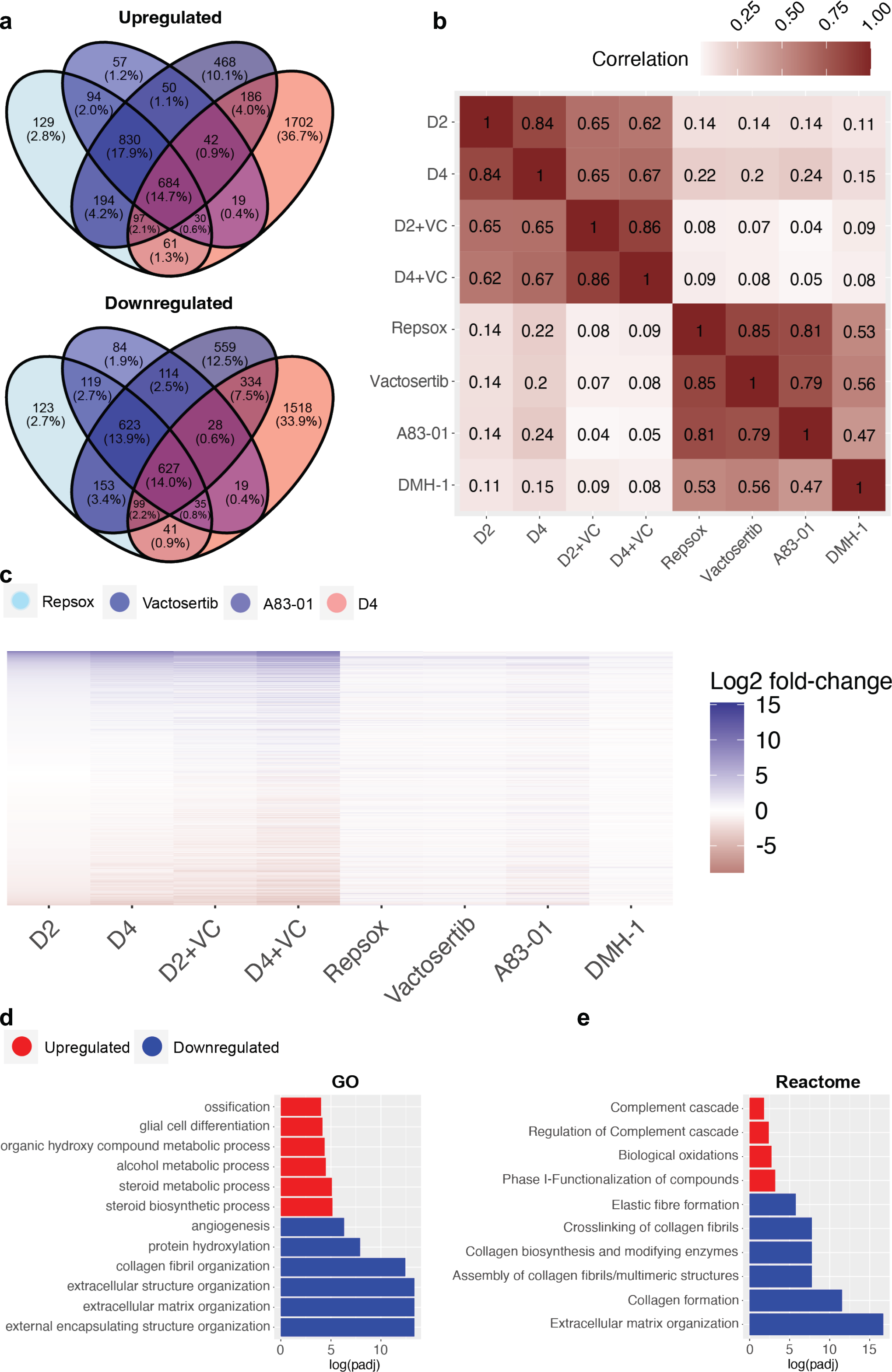
Transcriptomic comparison of Initation phase reprogramming with TGFb inhibition in Ercc1^Δ/−^. **(a)** Venn Diagram showing the overlap of significant genes (adjusted p-value < 0.05) for ALK5 inhibitors and 4 days of reprogramming. **(b)** Pairwise correlation between the log2 fold-changes of the filtered gene set (n=16883) of Ercc1 reprogramming and ALK5 or ALK2 inhibitors. **(c)** Heat map comparing Ercc1 reprogramming with TGFb inhibitors. Genes with a p-value lower than 1e-10 in at least one condition are shown. **(d and e)** GO and Reactome analysis of the intersect of significant genes as evaluated in 5A. Terms with the top six upregulated (red) and downregulated (blue) assessed by log(adjusted p-value) are shown.

Interestingly, significant differences were noted when comparing the transcriptomic effects of reprogramming vs TGFb inhibition on DNA repair processes even though both improved the DNA damage phenotype. Specifically, TGFb inhibition significantly downregulated multiple DNA repair processes in Ercc1 fibroblasts including AltEJ, NHEJ, and HR among others while reprogramming significantly upregulated them (Fig. S5a 3b). At the same time, only DMH-1 matched reprogramming by significantly upregulating NER, while Repsox and Vactosertib showed a trend towards improvment (Fig. s5a). In sum, this transcriptomic data demonstrates that TGFb inhibition and initiation phase reprogramming share a strong correlation in changes to gene expression. At the same time, reprogramming induces a more robust effect, especially on DNA repair processes in our reprogrammable Ercc1 aging model.

## Discussion

Although the age ameliorating effects of in vivo partial reprogramming have been well documented, the mechanistic drivers of rejuvenation by reprogramming are currently lacking^18^. Interestingly, the robust alterations to biological processes that occurs during the initiation phase of cellular reprogramming including proliferation, chromatin modification, DNA damage repair, MET, and RNA processing are well characterized and could potentially represent mechanistic drivers of longevity^41–45, 48^. For these reasons, we investigated the relationship between these early reprogramming induced changes and their impact on aging phenotypes. Specifically, we developed a reprogrammable DNA damage model of accelerated aging, Ercc1^Δ/−^ 4Fj^+/−^ rtTA^+/−^, in order to test the impact of initiation phase reprogramming on a key driver of aging, genomic instability. As OSKM inhibits TGFb signaling and drives the MET during initiation phase reprogramming, we investigated the mechanistic basis of rejuvenation via cellular reprogramming by comparing it to TGFb inhibition^46^. Due to the slow, stochastic, and asynchronous reprogramming process which confounds molecular studies in bulk cultures we also utilized a previously defined enhanced method of cellular reprogramming by adding ascorbic acid and CHIR-99021 to the culture media^53, 56, 57^.

Here we observed that OSKM expressing Ercc1 cells undergo robust reprogramming induced changes to morphology, nuclear size, epigenome, transcriptome, and homeostasis related DNA repair processes. Striking improvements to the DNA damage marker yH2AX in our Ercc1 fibroblasts with a defective repair process occurred within 2 days of pluripotency factor expression. At the same time, the elevated heterochromatin markers H3K9me3 and H4K20me3 were significantly restored following enhanced reprogramming for 2 and 4 days. Most notably, DNA methylation clock analysis shows significant rejuvenation in these Ercc1 cells after 4 days of enhanced reprogramming. RNA sequencing analysis of the transcriptome showed several GO processes were upregulated significantly including DNA repair, chromatin organization, and mitotic cell cycle. This is consistent with pluripotency transcription factors being upstream of DNA repair processes, particularly DNA double-strand break repair that repairs damage marked by yH2AX^76^. Simultaneously, TGFb signaling and EMT processes were significantly downregulated.

As TGFb signaling is a driver of many aging phenotypes, we further investigated the importance of this signaling pathway to reprogramming induced rejuvenation via inhibition of the TGFb receptor ALK5 or the BMP receptor ALK 2^67, 70^. Interestingly, nearly every TGFb inhibitor tested significantly decreased yH2AX levels in Ercc1 fibroblasts. The ALK5 inhibitors showed a more robust impact than ALK2 inhibitors on yH2AX levels while also decreasing nuclear size, a common phenotype observed during reprogramming. Importantly, rejuvenation of the epigenetic clock was observed following treatment with 3 of 4 TGFb inhibitors in this accelerated aging model. At the same time, RNA analysis showed a strong correlation between TGFb inhibition and reprogramming based on similar changes to hundreds of DEGs. Interestingly, the robust changes to DNA repair based on GO pathway analysis was unique to OSKM expression. Previous work has shown that activation of the TFGb pathway promotes genomic instability and induces HR defects while inhibition reduces DNA damage, senescence, and aging phenotypes^67, 74, 77, 78^. Interestingly, TGFb inhibition has a disruptive effect on many DNA repair processes while selectively promoting others in a cell type specific manner similar to our observations^65, 75^.

Although initiation phase reprogramming in wildtype and Ercc1 fibroblasts was capable of ameliorating the DNA damage marker yH2AX, the scale of effect was far more robust in the DNA damage model as demonstrated by significant increases in all major DNA repair processes. This data suggests that cellular reprogramming is sensitive to intrinsic cell state cues and drives a restoration of homeostatic function and youthful molecular phenotypes. Similarly, in vivo partial reprogramming studies show a more robust improvement to lifespan in progeria models indicating an ability to respond to deleterious cell states^23, 25^ Previously, it was shown improved DNA repair via overexpression of the HR protein Rad51 enhances reprogramming efficiency^79^. In our 4F Ercc1 fibroblasts, we observed the most significant reprogramming-induced rejuvenation coincided with upregulation of multiple DNA repair pathways. Recently maturation phase transient reprogramming for 13 days was shown to reverse the epigenetic clock in aged donors by up to 30 years but longer periods saw diminished benefits^33^. Other groups have shown a single short burst of reprogramming in vivo is sufficient to ameliorate aging hallmarks and tissue-specific physiology^25^. The precise mechanism of rejuvenation involved is unknown although several studies, including ours, supports a mechanistic contribution from the epigenetic remodeling process^6, 7, 32, 80^. Importantly, by inducing initiation phase reprogramming in an Ercc1 model with an accelerated epigenetic clock, we observed early reprogramming coincided with TGFb inhibition, improved homeostasis, DNA repair, and epigenetic rejuvenation. Notably, attempts to replicate OSKM-induced TGFb inhibition with small molecule inhibitors phenocopied improvements to the epigenetic clock in this Ercc1 model while decreasing the DNA damage marker yH2AX. Transcriptomic comparisons between cellular reprogramming and TGFb inhibitors of the ALK5 and ALK2 receptors were positively correlated and shared similar changes to hundreds of DEGs. At the same time, the robust upregulation of DNA repair processes via initiation phase reprogramming compared to TGFb inhibition alone suggests a different mechanistic route to DNA damage repair.

In conclusion, delineating the specific basis of rejuvenation remains difficult and potentially confounded by the multifactorial sequence of events necessary for reprogramming to proceed. The data here supports the premise that TGFb inhibition is a mechanistic contributor to rejuvenation induced by cellular reprogramming. This work sets the stage for further mechanistic investigations into the early events of reprogramming to better understand the associated drivers of rejuvenated phenotypes.

### Author contributions

A.O. and P.T.P. designed the study. A.O., A.H., R.B., S.H. and P.T.P. were involved in all experiments, data collection, analysis and interpretation. A.P and P.T.P generated the mice strains. P.T.P. and C.R. prepared the figures. C.M. performed skin fibroblast extractions. C.R., F.M., A.S., V.G. and P.T.P. prepared and analyzed RNA-seq data. A.O., P.T.P., and G.M. provided guidance. P.T.P wrote the manuscript with input from authors.

### Data availability statement

The data can be made available from the corresponding author upon request.

The mammalian methylation array is from the nonprofit Epigenetic Clock Development Foundation (https://clockfoundation.org/).

### Code availability statement

The RNA-seq analysis code is available at https://github.com/cheyennerechsteiner/Ercc1-Reprogramming.

## Methods

### Animal housing

These experiments were performed in accordance with Swiss legislation and after approval from the local authorities (Cantonal veterinary office, Canton de Vaud, Switzerland). Mice were housed in groups of five per cage or less with a 12hr light/dark cycle between 06:00 and 18:00 in a temperature-controlled environment at 25°C and humidity between 40 % and 70 %, with access to water and food. Wild type, premature aging, and programmable mouse models generated were housed together in the Animal Facilities of Epalinges and Department of Biomedical Science of the University of Lausanne.

### Mouse strains

Ercc1^Δ/−^ mice and littermate controls Ercc1^+/+^ were generated in a C57BL6J|FVB hybrid background as previously generated and described by de Waard ^13^. Reprogrammable 4Fj^+/−^ rtTA^+/−^ Ercc1^Δ/−^ (4F.D/KO) and 4Fj^+/−^ rtTA^+/−^ Ercc1^+/+^ (4F.WT) mice strains were generated in a C57BL6J|FVB hybrid background following crosses with the previously described 4Fj mouse developed by Rudolf Jaenisch ^82^. Previously described wildtype reprogrammable cells with an Oct4-EGFP reporter were derived from mice carrying a polycistronic cassette containing Oct4, Sox2, Klf4, c-Myc, and a single allele EGFP in the Col1a1 locus (OKSM)^83^.

### Mouse monitoring and euthanasia

All mice were monitored three times per week for their activity, posture, alertness, body weight, presence of tumors or wound, and surface temperature. Males and females were euthanized at by CO2 inhalation (6 min, flow rate 20% volume/min). Subsequently, tail tip fibroblasts were harvested and cultured.

### Cell culture and maintenance

Tail tip fibroblasts (TTF) were extracted from mice using Collagenase I (Sigma, C0130) and Dispase II (Sigma, D4693) and cultured in DMEM (Gibco, 11960085) containing 1% non-essential amino acids (Gibco, 11140035), 1% GlutaMax (Gibco, 35050061), 1% Sodium Pyruvate (Gibco, 11360039) and 10% fetal bovine serum (FBS, Hyclone, SH30088.03) at 37°C in hypoxic conditions (3% O2). Subsequently, fibroblasts were cultured and passaged (3 or less) according to standard protocols. Reprogrammable TTF cells used in experiments were from young mice, including 4Fj^+/−^ rtTA^+/−^ Ercc1^Δ/−^ and 4Fj^+/−^ rtTA^+/−^ Ercc1^+/+^ (8 week old male), and 4FJ^+/-^ OCT4-GFP TTF (22 week old female). Non-reprogramming Ercc1 TTF used include Ercc1^Δ/−^ and Ercc1^+/+^ (8 week old male) Experiments for TGFb inhibitors used Ercc1^Δ/−^ cells (4 week old female) and Ercc1^+/+^ TTF cells (14 week old female). Induction of the OSKM reprogramming factors in vitro followed treatment with doxycycline 2ug/ml in culture media for the specified time points. Enhanced reprogramming followed the same protocols but added fresh ascorbic acid (50ug/ml) and CHIR-99201 (3uM) as previously reported^53^.

### Immunofluorescence staining

Following experiments in 96 well plates or 24 well plates, TTF were washed with fresh PBS and fixed with 4% paraformaldehyde (Roth, 0964.1) in PBS at room temperature (RT) for 15 minutes. Then cells were washed 3 more times, followed by blocking and permeabilization step in 1% bovine serum albumin (Sigma, A9647-50G) in PBST (0.2% Triton X-100 in PBS) for 1 hour (Roth, 3051.3). TTF were then incubated at 4°C overnight with a primary antibody, washed in PBS, followed by incubation with a secondary antibody and DAPI staining at room temp. for 1 hour. If performed in 24 well plates, coverslips were used and mounted with Fluoromount-G (Thermofisher, 00-4958-02), dried at RT in the dark, and stored at 4°C until ready to image or -20°C for long-term.

### Immunofluorescence imaging

Confocal image acquisition was performed using the NIKON Ti2 Yokogawa CSU-W1 Spinning Disk, using the 100X objective and with 15 z-sections of 0.3 μm intervals. Lasers for each antibody were selected (405 nm and 488 nm) with a typical laser intensity set. Exposure time and binning were established separately to assure avoidance of signal saturation.

### Antibodies and compounds

Antibodies include: Abcam: anti-H3K27me3 (ab192985); Cell Signaling: anti-H3K9me3 (13969), anti-H4K20me3, anti-γH2AX (9718); Roth: DAPI (6843.1) Compounds include: Cayman; Valproic Acid (13033), CHIR99021 (13122), Repsox (14794), Forskolin (11018), Doxorubicin (15007); Acros Organics: TCP (130472500); APExBIO: DZNep (A8182); Seleckchem: TTNPB (S4627); Roth: X-beta-Gal (2315.3), Ascorbic Acid (Sigma).

### Flow cytometry

Cells were stained with Thy1.2-APC and SSEA1-PE (Biolegend). Cells were fixed in BD Fixation and Permeabilization Solution. A Bechman Coulter Cytoflex S flow cytometer determined cellular subpopulation ratios.

### RNA sequencing, processing, analysis

Total RNA and DNA was extracted from the same samples. Total RNA was extracted from cells using the Qiagen AllPrep DNA/RNA Micro Kit (New England Biolab) and protocols were followed. Total RNA concentrations were determined using the Qubit RNA BR Assay Kit (Thermofisher, Q10211).

The RNA-Seq library preparation and sequencing was done by Novogene (UK) Company Limited on an Illumina NovaSeq 6000 in 150 bp paired-end mode. Raw FASTQ files were evaluated for quality, adapter content and duplication rates with FastQC. Reads were trimmed using TrimGalore! (https://www.bioinformatics.babraham.ac.uk/projects/trim_galore/) and aligned to the GRCm39 mouse genome assembly using Hisat2 (https://doi.org/10.1038/s41587-019-0201-4). Number of reads per gene were measured using the featureCounts function in the subread package^84^.

All subsequent analysis was performed in R. DESeq2 was used to normalize raw read counts and perform differential gene expression analysis (doi:10.1186/s13059-014-0550-8). Ensembl gene IDs were mapped to gene symbols via the mapIds function in the AnnotationDbi package (Pagès H, Carlson M, Falcon S, Li N (2023). *AnnotationDbi: Manipulation of SQLite-based annotations in Bioconductor*) with an org.Mm.eg.db reference package (Carlson M (2019). *org.Mm.eg.db: Genome wide annotation for Mouse*). The clusterProfiler package (doi:10.1089/omi.2011.0118, doi:10.1016/j.xinn.2021.100141) was used to perform gene set enrichment analysis (GSEA) on gene ontology and Reactome terms.

### DNA extractions

Total DNA was extracted from cells using the Qiagen AllPrep DNA/RNA Micro Kit (New England Biolab) and protocols were followed. Total DNA concentrations were determined with the Qubit DNA BR Assay (Thermofisher, Q10211).

### DNA methylation clock

Methylation data was generated via the HorvathMammalMethylChip^85^ and normalized with the SeSaMe method^86^. Human methylation data were generated on the Illumina EPIC array platforms which profiles 866k cytosines. The noob normalization method was used and implemented in the R function preprocess Noob. The DNAm age was estimated using the DNAm Age Skin Final clock algorithm.^87^

### MTS cell proliferation assay

Cell viability was performed with Tetrazolium MTS assay. Cells were cultured for 1 day in 96-well plates and treated with small molecules for 3 consecutive days prior to incubation with 120 μL fresh media containing 20 μL of CellTiter 96® AQueous One Solution (Promega, G3580) for 1 to 4 hours at 37°C in a humidified, 5% CO2 atmosphere. The absorbance was determined at 490nm using a BioTek Epoch 2 microplate reader. The proportion of viable cells was determined as a ratio between the observed optical density (OD) compared to control OD.

### Quantification and statistical analysis

Analysis of immunofluorescence microscopy images was performed using FIJI. Typically, 20 different ROIs were imaged at 100x from each well from three experiments unless otherwise noted. Maximal projections of z-stacks were analyzed and total fluorescence intensity per cell and total nuclear area were determined.

All statistical analysis, statistical significance and n values are reported in the figure legends. Statistical analyses were performed using GraphPad Prism 9.0.0.

---

**Figure S1:**
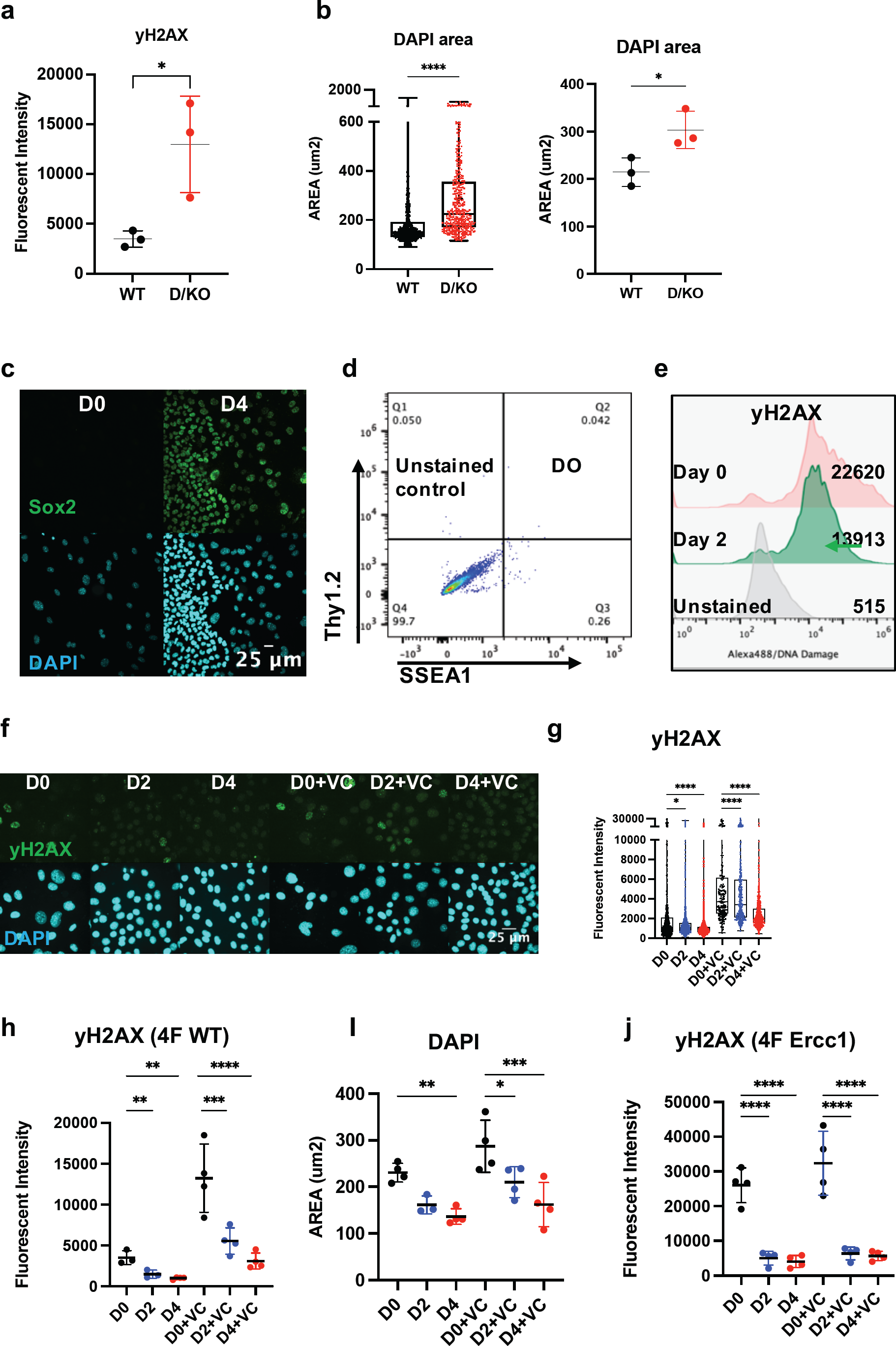
Initiation phase reprogramming in 4Fj^+/−^ rtTA^+/−^ Ercc1^Δ/−^ accelerated aging model promotes DNA damage repair. **(a)** Quantification of yH2AX IF from 3 experiments in fibroblasts from wildtype (WT) and Ercc1^Δ/−^ (D/KO) mice, imaged with Nikon laser confocal spinning disc, according to unpaired t-test, * p<0.05. **(b)**. Quantification of DAPI in fibroblasts from wildtype (WT) and Ercc1^Δ/−^ (D/KO) mice, imaged with Nikon laser confocal spinning disc, according to Mann-Whitney test, **** p<0.0001. Quantification of DAPI area from 3 experiments in fibroblasts from wildtype (WT) and Ercc1^Δ/−^ (D/KO) mice, imaged with Nikon laser confocal spinning disc, according to unpaired t-test, * p<0.05 **(c)** IF image of Sox2 and DAPI in 4F.D/KO Fbs after 4 days doxycycline induction, 40x **(d)** FACS unstained control for analysis of reprogramming Fbs markers Thy1.2 and SSEA1. **(e)** FACS analysis yH2AX levels following reprogramming of 4F.D/KO Fbs at day 0, day 2, and unstained control. **(f)** IF images of yH2AX and DAPI during representative time course analysis of 4F.WT Fb +/-VC at 100x. **(g)** Quantification of yH2AX images during time course shows significantly decreased levels at day 2 and day 4 according to one way ANOVA, **** p<0.0001. **(h)** Quantification of mean yH2AX levels in 4F WT from 3 or 4 experiments during time course shows significant decrease to levels at day 2 and day 4 according to one way ANOVA, ** p<0.01. **(i)** Quantification of DAPI area during time course shows decrease in size at day 2 and day 4 based on mean values of 4 experiments according to one way ANOVA, *** p<0.001, ** p<0.01, *p<0.05. **(j)** Quantification of mean yH2AX levels in 4F D/KO from 4 experiments during time course shows significant decrease to levels at day 2 and day 4 according to one way ANOVA, ** p<0.0001.

**Figure S2:**
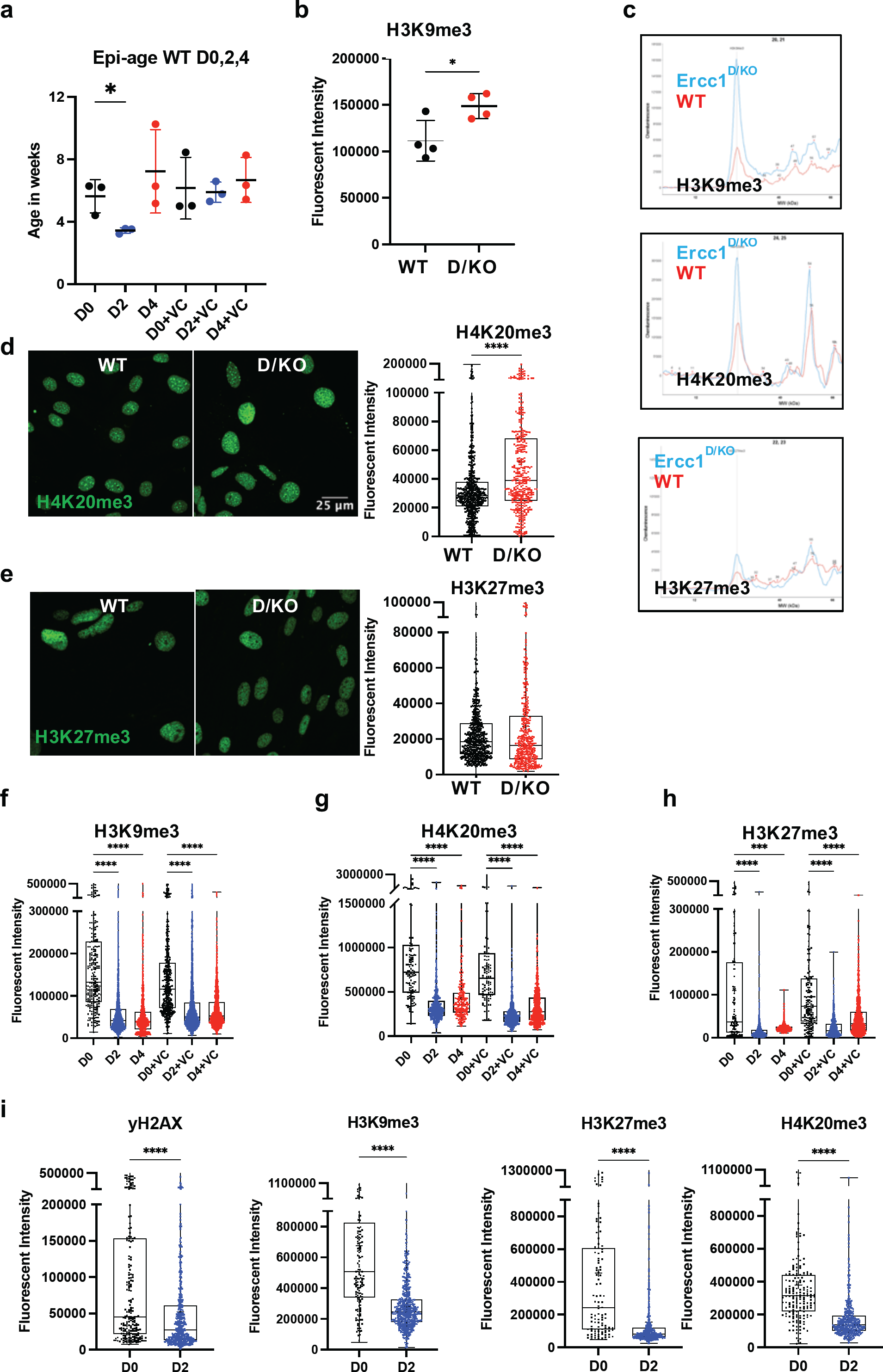
DNA methylation clock is restored in Ercc1^Δ/−^ following short term reprogramming. **(a)** DNA Methylation Age Skin Final clock analysis of WT Fb during time course, n = 3, * p<0.05 according to unpaired t-test. **(b)** Quantification of mean values of H3K9me3 levels in fibroblasts from WT and D/KO mice (n=4) according to unpaired t-test, * p<0.05. (**c)** Western capillary chemiluminescence quantification shows increased H3K9me3 and H4K20me3, with unchanged H3K27me3 in fibroblasts from D/KO vs WT mice (n=1 each). **(d)** IF and quantification shows increased H4K20me3 in fibroblasts from D/KO mice vs WT, imaged with Nikon laser confocal spinning disc, according to Mann-Whitney test, **** p<0.0001 **(e)** IF quantification shows unchaged H3K27me3 in fibroblasts from WT and D/KO mice, imaged with Nikon laser confocal spinning disc. **(f, g,h)** IF quantification of 2^nd^ experiment shows significantly decreased H3K9me3, H4K20me3, and H3K27me3 levels during time course in 4F.D/KO Fbs after 2 and 4 days doxycycline induction with or without VC at 100x according to Kruskal-Wallis test, * p<0.000. **(i)** IF quantification of 3^rd^ experiment shows yH2AX, H3K9me3, H4K20me3, and H3K27me3 in 4F.D/KO Fbs after 2 doxycycline induction at 100x according to Mann-Whitney test, * p<0.000.

**Figure S3:**
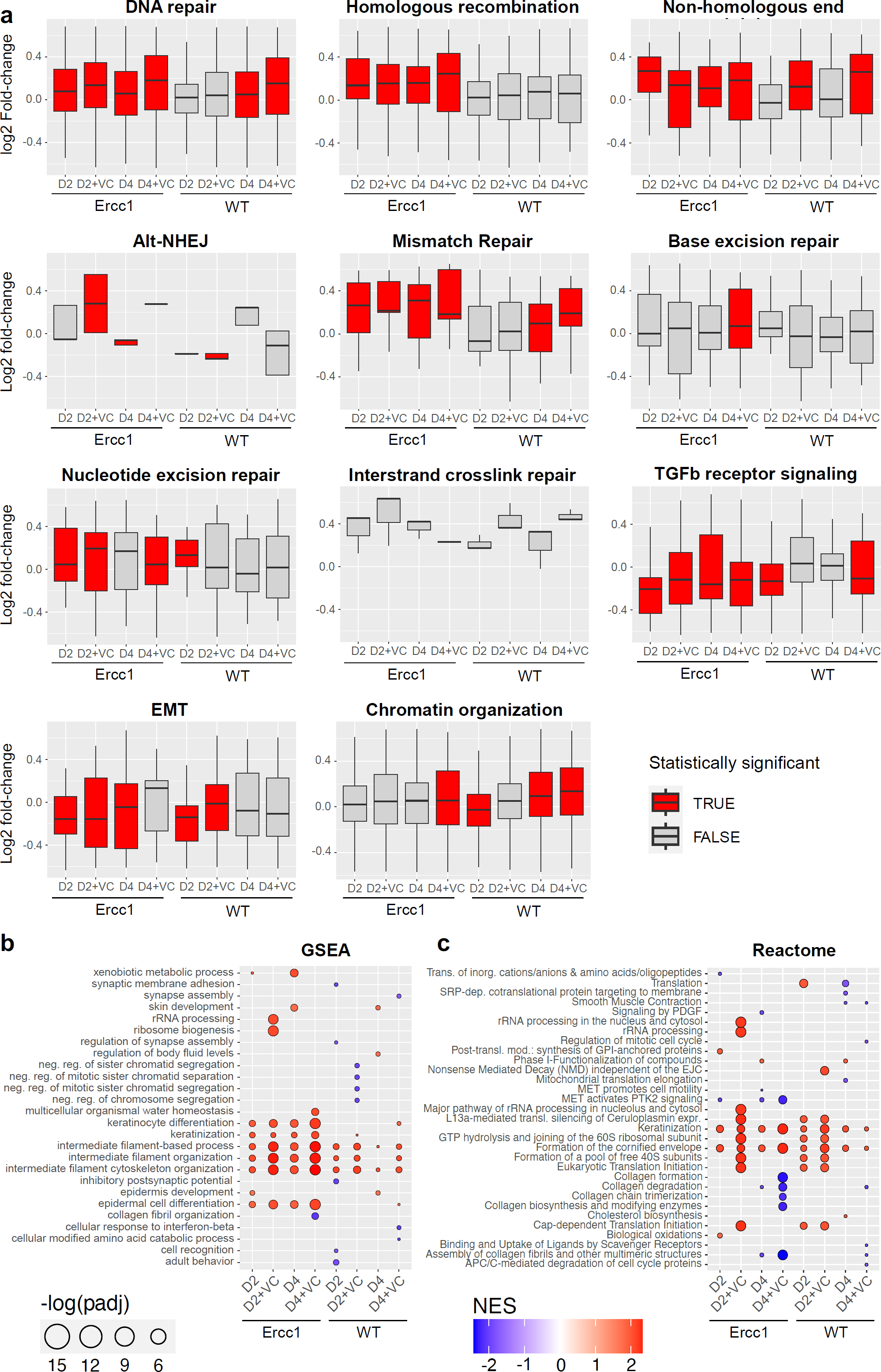
Initation phase reprogramming drives transcriptomic reset and upregulates DNA repair and chromatin organization in Ercc1^Δ/−^. **(a)** Log2 fold-change of genes within a given GO pathway for Ercc1 and WT reprogramming at days 0, 2, and 4 with and without VC enhancement. The inner boxplot depicts medians and the first and third quartiles, with whiskers extending up to the 1.5x interquartile range and outliers removed for improved visualization of differences between conditions. Statistical significance (Wilcoxon Test, p-value < 0.05) is indicated by red coloring. **(b and c)** GSEA and Reactome analysis of Ercc1 and WT reprogramming considering log2 fold-changes between day 0 and day 2 of reprogramming and day 0 and day 4 of reprogramming with and without VC enhancement. Gene ontology biological process terms and Reactome terms are plotted against the normalized enrichment score (NES) with -log(adjusted p-value) illustrated through circle size. The top 10 terms for each condition are included.

**Figure S4:**
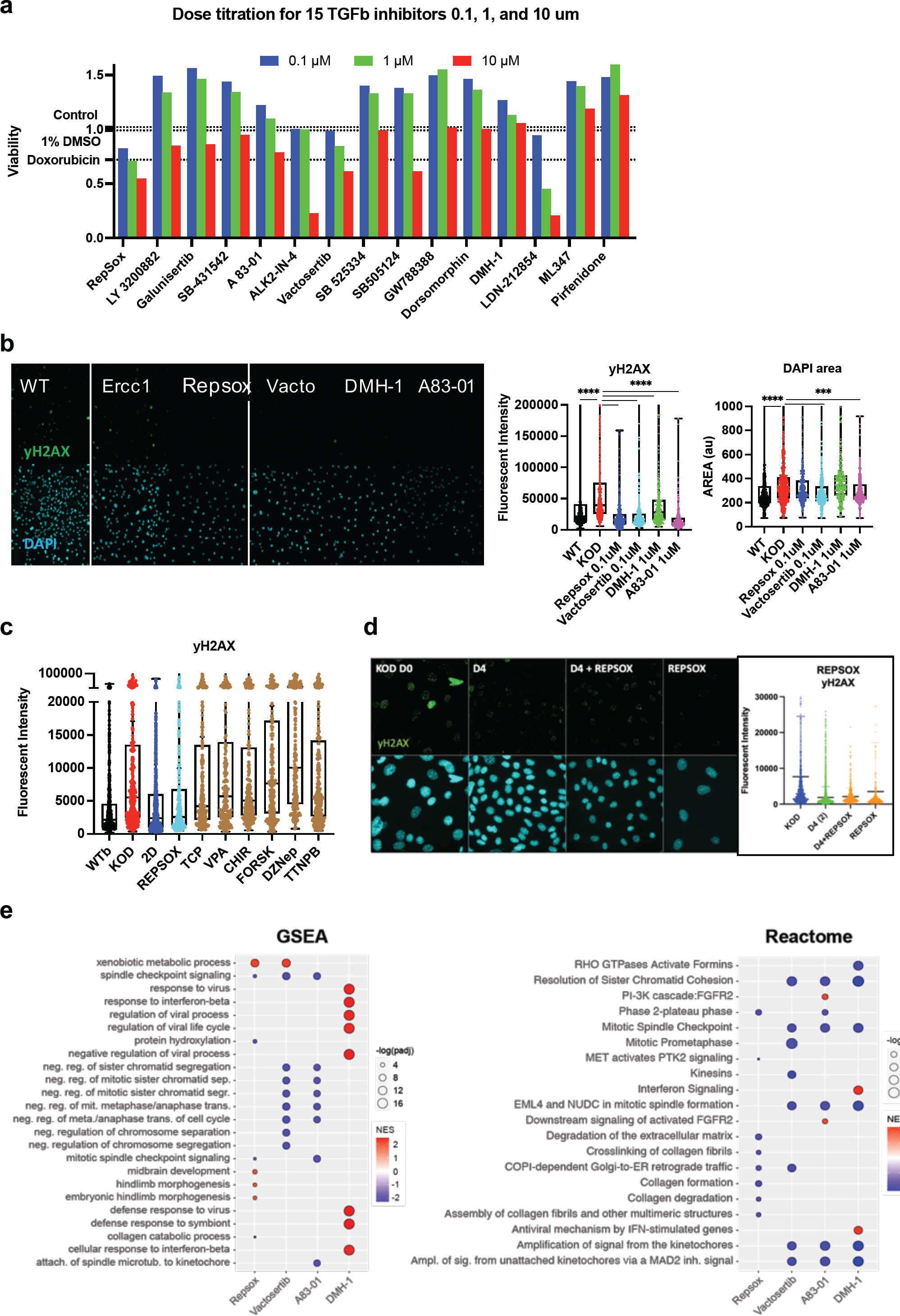
Inhibition of ALK5 or ALK2 receptors improves DNA damage phenotype and resets the DNA methylation clock and transcriptome in Ercc1^Δ/−^. **(a)** Cell viability quantification using MTS assay in 96 well plates following treatment with high, medium, and low concentrations of 15 inhibitors in triplicate. Dosage range is 10um, 1um, and 0.1 um. Controls are represented by dashed lines and include untreated D/KO Fbs, 1% DMSO, and Doxorubicin (0.1um). **(b)** IF images and quantification of yH2AX levels and DAPI area in WT vs D/KO Fbs treated with Repsox, Vactosertib, DMH-1, and A83-01 in 24 wells plates, n = 1. **(c)** IF quantification of yH2AX levels comparing 2 days of doxycline induction to seven single small molecules used in chemical reprogramming at published dosages: Repsox (3um), TCP (10um), VPA (500um), CHIR99021 (5um), Forskolin (10um), DZNep (0.5um), and TTNPB (1um), n = 1. **(d)** IF images and quantification of yH2AX levels in 4F.D/KO cells after 4 days of induction either with doxycycline alone, doxycycline and Repsox, or Repsox alone, n = 1. **(e)** GSEA and Reactome analysis of Ercc1 cells treated with TGFb inhibitors considering log2 fold-changes, n = 3. Gene ontology biological process terms or Reactome terms are plotted against the normalized enrichment score (NES) with -log(adjusted p-value) indicated through circle size. The top 10 terms for each condition are included.

**Figure S5:**
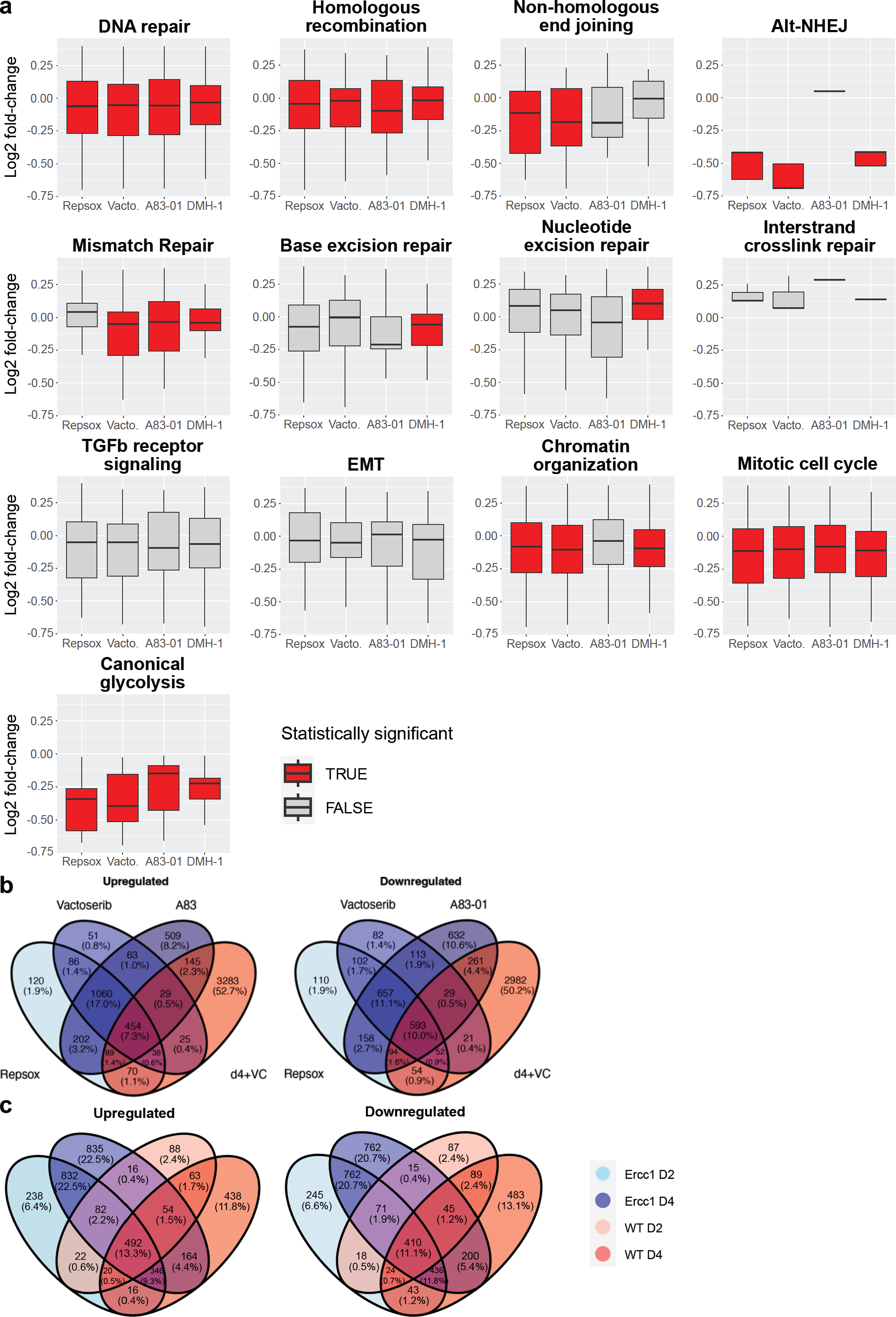
GO term analysis of TGFb inhibition in Ercc1^Δ/−^. **(a)** Log2 fold-change of genes within a given GO pathway for Ercc1 cells treated with TGFb inhibitors compared to non-treated Ercc1 cells. The inner boxplot depicts medians and the first and third quartiles, with whiskers extending up to the 1.5x interquartile range and outliers removed for improved visualization of differences between conditions. Statistical significance (Wilcoxon Test, p-value < 0.05) is indicated by red coloring. **(a)** Venn Diagram showing the overlap of significant genes (adjusted p-value < 0.05) for the ALK5 inhibitors and enhanced reprogramming day 4+VC. **(c)** Venn Diagram showing the overlap of significant genes (adjusted p-value < 0.05) for the reprogramming time course at day 2 and 4 without VC.

